# Allele-Specific Effects of Mutations in the Rifampin Resistance-Determining Region (RRDR) of RpoB on Physiology and Antibiotic Resistance in *Enterococcus faecium*

**DOI:** 10.1101/2025.07.25.666922

**Authors:** Adeline Supandy, Emma G. Mills, Kyong T. Fam, Ryan K. Shields, Howard C. Hang, Daria Van Tyne

## Abstract

*Enterococcus faecium* is a member of the human gut microbiota that has evolved into a problematic nosocomial pathogen and leading cause of infections in hospitalized patients. Treatment of *E. faecium* infections is complicated by antibiotic resistance, making it important to understand resistance mechanisms and their broader consequences in this pathogen. Here we explored the collateral effects of rifampin resistance-associated mutations in the *E. faecium* RNA polymerase β-subunit (RpoB). Of 14,384 publicly available *E. faecium* genomes, nearly one-third carried a mutation in the rifampin resistance-determining region (RRDR) of RpoB. In a local population of 710 *E. faecium* clinical isolates collected from patients at a single hospital, we found significant associations between the presence of RRDR mutations and prior exposure to rifamycin antibiotics, as well as associations between RRDR mutations and altered daptomycin susceptibility. To investigate the phenotypic impacts of RRDR mutations, we generated and studied four isogenic strains with distinct RRDR mutations (Q473K, G482D, H486Y, S491L) that overlapped with clinical isolate variants. Transcriptomic and phenotypic analyses revealed allele-specific effects on *E. faecium* gene expression, growth dynamics, antibiotic susceptibility, isopropanol tolerance, and cell wall physiology. One frequently observed mutation, H486Y, caused minimal transcriptional changes and enhanced bacterial fitness under antibiotic stress. In contrast, the S491L mutation induced extensive transcriptional changes and slowed bacterial growth, but also conferred increased isopropanol tolerance, potentially enhancing bacterial survival in the hospital. Overall, our findings highlight the multifaceted impacts of RRDR mutations in shaping *E. faecium* physiology and antibiotic resistance, two important features of this hospital-associated pathogen.

**Importance:** Understanding how antimicrobial resistance affects bacterial physiology is critical for developing effective therapeutics against bacterial infections. In this study, we found that rifampin resistance-associated mutations in RpoB are widespread in *Enterococcus faecium,* a leading multidrug-resistant pathogen. By studying isogenic wild-type and RpoB mutant strains, we discovered that RpoB mutations, while conferring resistance to rifampin, have distinct allele-specific effects on other bacterial phenotypes. Some of these collateral effects appear to promote *E. faecium* resistance to antibiotics and survival in the hospital environment, raising questions about the selective pressures driving their emergence. Overall, our study underscores the importance of examining the collateral effects of resistance-associated mutations in multidrug-resistant pathogens, which could help mitigate their persistence and spread among vulnerable patients.

## Introduction

Antimicrobial resistance represents one of the most pressing public health challenges in the 21^st^ century, with nearly 5 million deaths attributed to resistant infections each year^1,2^. Among the pathogens driving this crisis, *Enterococcus faecium* causes invasive infections that are often antibiotic-resistant and difficult to treat. The ability to rapidly adapt to diverse environments allows *E. faecium* to overcome bottlenecks caused by selective pressures such as antibiotic exposure, nutrient limitation, and host immunity. As such, among the ESKAPE pathogens, *E. faecium* is a leading cause of bloodstream infections (BSIs), especially in hospitalized and immunocompromised patients^3,4^. In addition to various intrinsic resistance mechanisms, *E. faecium* also possesses the ability to rapidly acquire additional resistance genes^5–7^. Treatment is particularly challenging due to the organism’s ability to resist several important gram positive-targeting antibiotics, including vancomycin, ampicillin, and daptomycin^6–11^. Consequently, *E. faecium* is often associated with treatment failure and recurrent infections, contributing to an estimated 100,000 deaths globally each year^12^. Understanding the mechanisms underlying this adaptability is crucial to developing new therapeutics against *E. faecium* infections.

While the genetic determinants of antibiotic resistance in *E. faecium* have been studied extensively, many resistance mechanisms have yet to be fully elucidated^5^. In particular, recent studies have demonstrated that mutations in the DNA-dependent RNA polymerase β-subunit (RpoB), particularly in the 81-bp Rifampin Resistance-Determining Region (RRDR), may play a broader role beyond rifampin resistance^13,14^. For example, RRDR mutations have been associated with increased resistance to cephalosporins and daptomycin, the latter of which is a last-resort antibiotic for *E. faecium* infections^13,14^. These findings suggest that RRDR mutations may have pleiotropic effects on antibiotic susceptibility and resistance in *E. faecium*. Additionally, it is unclear whether the emergence of RRDR mutations in *E. faecium* is driven solely by exposure to rifamycin class antibiotics, or whether other environmental factors might contribute to their selection. Because RpoB is an essential and highly conserved component of the bacterial transcriptional machinery, mutations in this protein are likely to affect a variety of cellular processes through altered gene expression^15^. A deeper understanding of how RRDR mutations emerge and their impact on *E. faecium* physiology is crucial to the development of new treatments for *E. faecium* infections.

In this study, we investigated the global prevalence, diversity, and functional impact of mutations in the RRDR of RpoB in *E. faecium*. We determined the global distribution of RRDR mutations and confirmed previously described associations between RRDR mutations and daptomycin susceptibility in a cohort of over 700 patients from a single hospital. We also used genomic, transcriptomic and phenotypic analyses to investigate allele-specific effects of RRDR mutations on *E. faecium* physiology and resistance to antibiotics beyond rifampin. Taken together, our findings highlight the multifaceted roles of RRDR mutations in shaping *E. faecium* biology.

## Results

### RRDR mutations are prevalent in human-associated *E. faecium*

To understand the global epidemiology of RRDR mutations in *E. faecium*, we first analyzed all publicly available genomes deposited in the National Center for Biotechnology Information (NCBI) database between 2000 and 2023. The dataset was filtered to include 14,384 human-associated *E. faecium* genomes, with most isolates originating from Europe (52%), North America (25%), and Oceania (16%) (Supplemental Figures 1 and 2, Supplemental File 1). Approximately 30% of genomes encoded one or more mutations in the RRDR (Figure 1A; Supplemental Figure 3). The most frequent mutation, S491F, was found in nearly 20% of all global *E. faecium* genomes and appeared to increase in frequency starting in 2007 (Figure 1B). This and the second most prevalent variant, the H486Y/M475V double mutant, were frequently found among isolates belonging to the emerging lineages sequence type (ST) 80 and ST117 (Figure 1C)^16^. Many RRDR mutations were found across multiple countries, STs, and genetic backgrounds, suggesting independent emergence.

**Figure 1.**
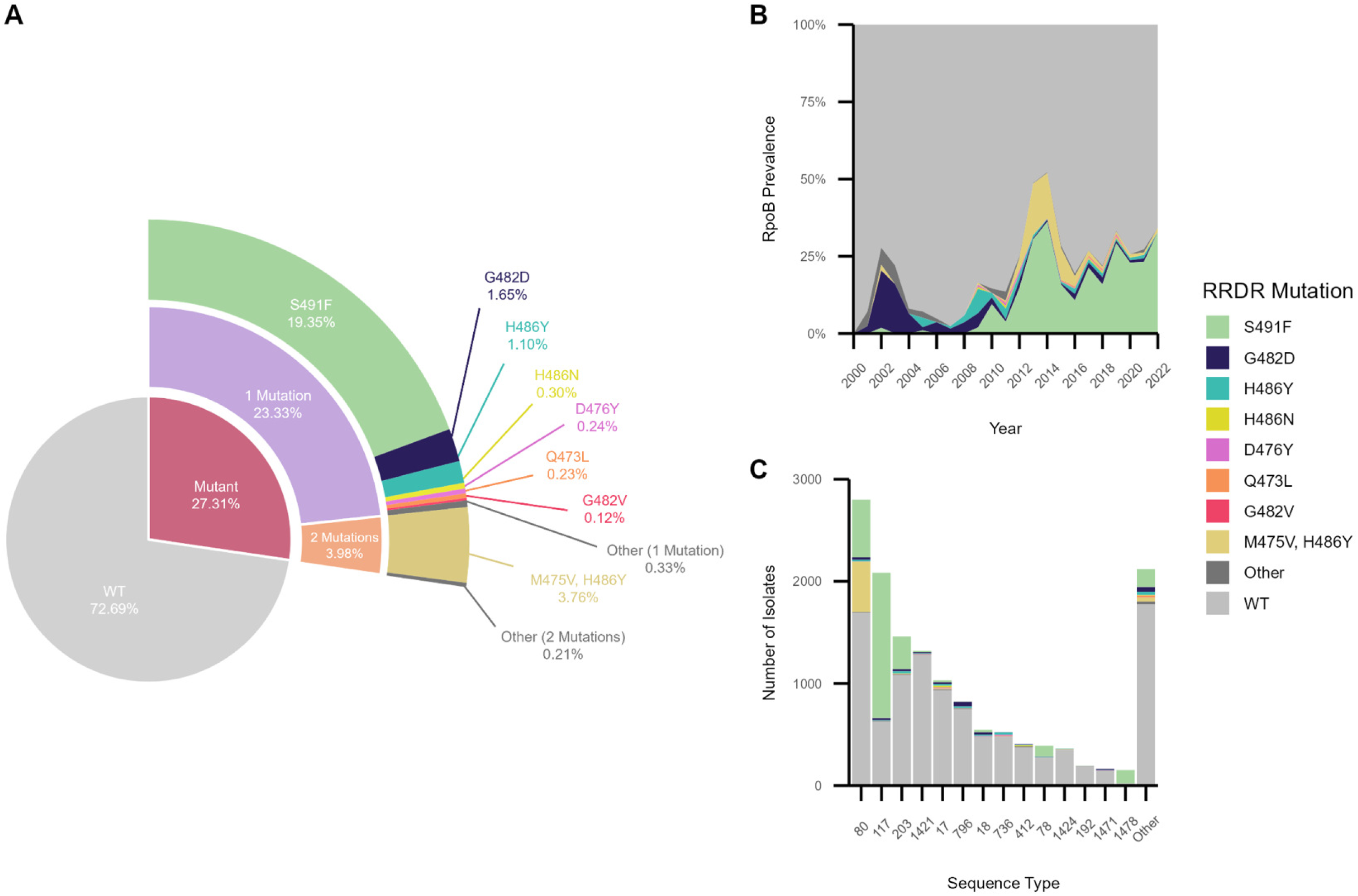
Distribution of mutations in the RpoB rifampin resistance-determining region (RRDR) across 14,384 publicly available *E. faecium* isolate genomes. RRDR variants were categorized based on total variant frequency (A), year of isolation (B), and sequence type (ST) (C). Data were collected from NCBI and filtered according to Supplemental Figure 1 to include only human-associated *E. faecium* collected between 2000 to 2023.

**Figure 2.**
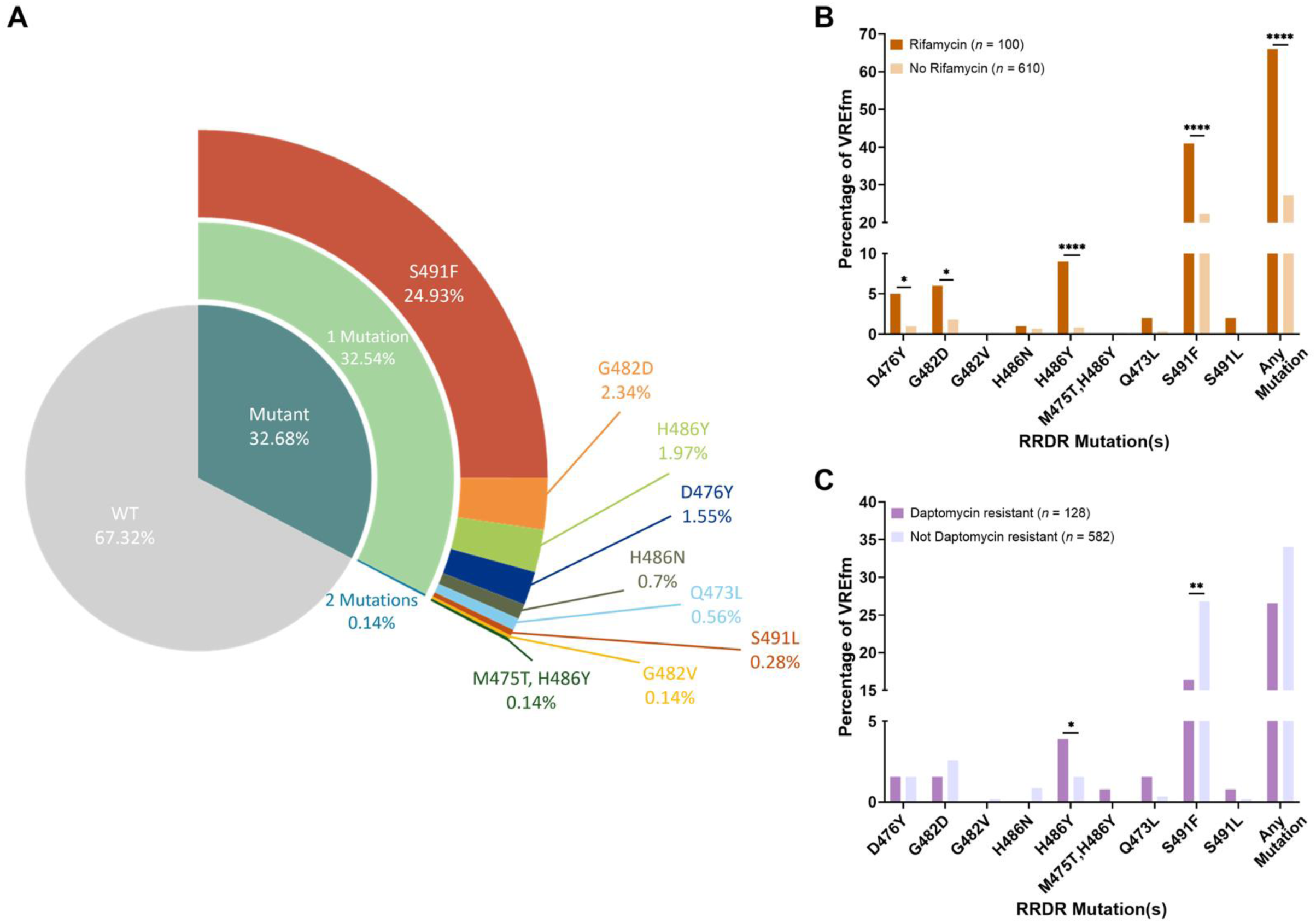
Associations between RRDR mutations and prior antibiotic exposure and resistance in 710 vancomycin-resistant *E. faecium* (VREfm) isolates collected from a single medical center. (A) Distribution of RRDR mutations across VREfm isolates based on total variant frequency. (B) Associations between the percentage of VREfm isolates with RRDR mutations in patients with prior exposure to antibiotics in the rifamycin class (*n* = 100) or patients without exposure (*n* = 610). (C) Associations between the percentage of VREfm isolates with RRDR mutations that are daptomycin-resistant (*n* = 128) or daptomycin-susceptible (*n* = 582). Data were analyzed with one-way ANOVA. *p≤0.05; **p≤0.01; ****p≤0.0001.

As RRDR mutations have been shown to confer rifampin resistance, we sought to assess the association between RRDR mutations and prior exposure to rifamycin class antibiotics among 710 *E. faecium* isolates collected from patients at the University of Pittsburgh Medical Center (UPMC) between 2017 and 2022^16–19^. Similar to the global distribution, roughly 33% of these isolates encoded RRDR mutations, with S491F being the most prevalent (25%) (Figures 1A and 2A). Most of the RRDR mutations detected were strongly associated with prior exposure to rifamycin class antibiotics, including the D476Y, G482D, H486Y, and S491F variants (Figure 2B). A prior study found that RRDR mutations were also associated with daptomycin resistance in *E. faecium*^14^, and we wondered whether similar associations were present in this dataset. While prior exposure to daptomycin was not associated with RRDR mutations, we observed that two mutations were significantly associated with altered daptomycin susceptibility (Figure 2C). The H486Y mutation was associated with increased daptomycin resistance, however the S491F mutation was associated with daptomycin sensitivity. While these results do not entirely agree with the prior study^14^, they nonetheless suggest that RRDR mutations are prevalent and could confer additional advantages that benefit *E. faecium* expansion in clinical settings.

### RRDR mutations induce widespread transcriptional changes in *E. faecium*

Since the RNA polymerase plays a critical role in gene transcription, we wondered how different RRDR mutations altered *E. faecium* transcriptional profiles at a genome-wide scale^20–24^. To investigate this question, we generated four isogenic RRDR mutants through one-step selection by plating a clinical *E. faecium* isolate onto media containing rifampin^25^. Whole-genome sequencing of individual clones confirmed that the resulting rifampin-resistant isolates were isogenic and carried G482D, Q473K, H486Y, or S491L mutations in the RRDR. The G482D and H486Y mutations were also observed in the human-associated *E. faecium* datasets, while mutations at residues Q473 and S491 were also detected, albeit with substitutions to different amino acids (Figure 1A and 2A).

To measure the effects of different RRDR mutations on *E. faecium* transcription, we performed RNA sequencing on the wild-type (WT) and isogenic mutant strains and identified differentially expressed genes in each mutant (Figure 3A-D). The overall number of differentially expressed genes in each strain varied widely, with many genes down-regulated in the Q473K, G482V, and S491L mutants and a small number of genes up-regulated in the H486Y mutant. Principal component analysis revealed tight clustering of replicate transcriptomes of each strain, however, transcriptomes of different strains clustered apart from one another (Figure 3E). H486Y mutant transcriptomes clustered closely with those of the WT strain, consistent with the lower number of differentially expressed genes in this strain (Figure 3E and 3F). We also compared the differentially expressed genes in the mutant strains to one another and found that many genes were unique to each mutant (Figure 3F; Supplemental Figure 4A-B). Ten genes were differentially expressed in all four mutant strains (Figure 3F); however, these genes were all down-regulated in the Q473K, G482V, and S491L mutants but up-regulated in the H486Y mutant (Supplemental File 1).

**Figure 3.**
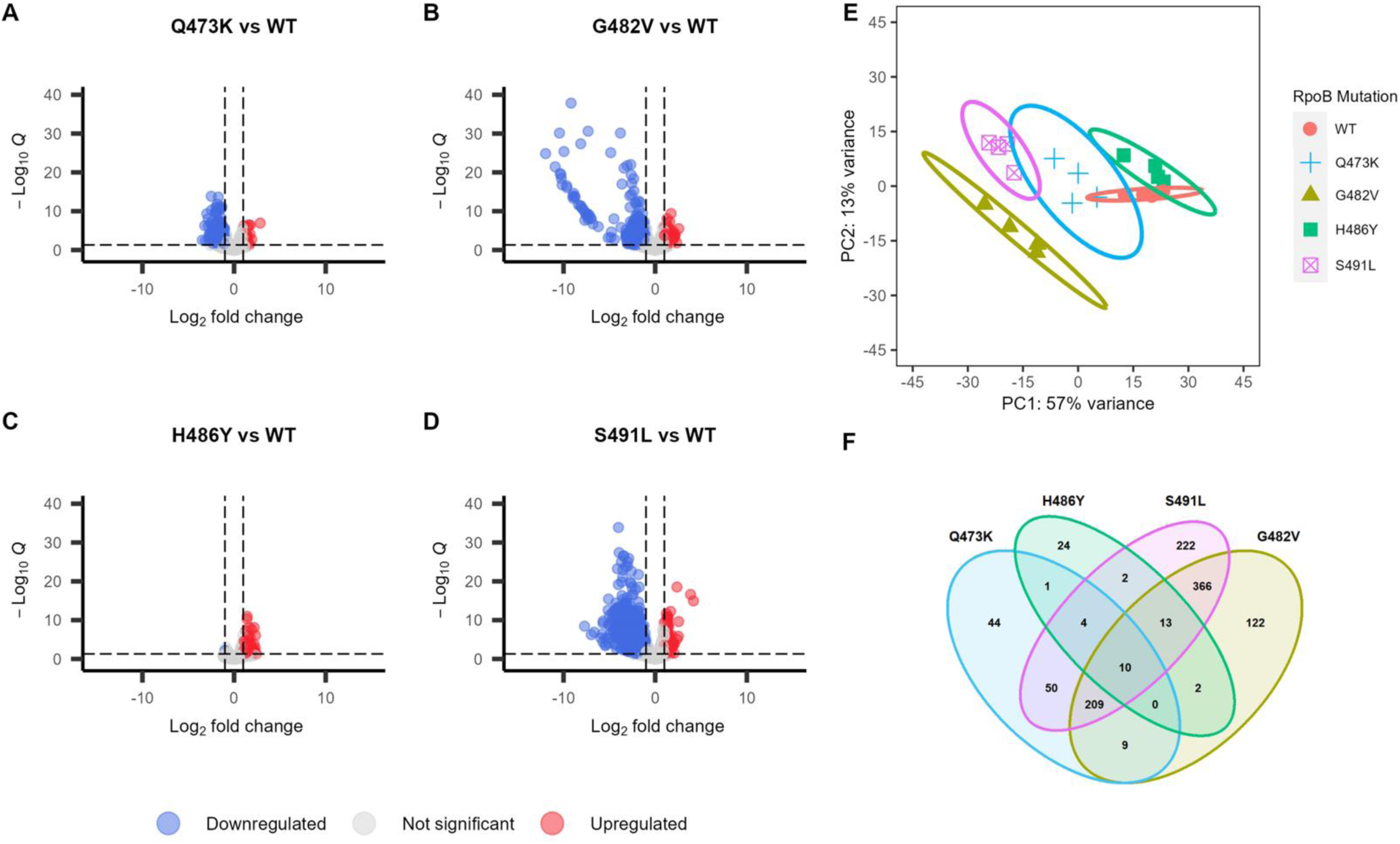
Visualization of differentially expressed genes (DEGs) due to mutations in RRDR as identified by RNA sequencing. (A - D) Volcano plots showing the distribution of up- and down-regulated genes in each RRDR mutant. The x-axis shows log_2_ fold-change in expression level and the y-axis shows the significance of the difference in expression. (E) Principal Component Analysis of RNA-seq samples with 95% confidence intervals circled. Different RRDR mutants are represented by different colors and shapes. Each dot represents one biological replicate. (F) Venn diagram showing the number of shared differentially expressed genes between RRDR mutants.

We next investigated the functional roles of differentially expressed genes in each mutant strain by analyzing their distribution of Clusters of Orthologous Groups (COG) categories relative to the distribution of all genes in the WT strain genome^26,27^. Among up-regulated genes, only COG category F (Nucleotide Transport and Metabolism) was significantly enriched in the H486Y mutant (Figure 4). The H486Y mutant had a single gene down-regulated, while the other three mutant strains all showed significant down-regulation of genes involved in Carbohydrate Transport and Metabolism (Category G; Figure 4). Both the S491L and G482V mutants also demonstrated significant down-regulation of genes related to Translation, Ribosome Structure, and Biogenesis (Category J; Figure 4). Together these findings suggest that RRDR mutations differentially affect global transcription and primarily target metabolic pathways, potentially impacting *E. faecium* fitness and growth dynamics.

**Figure 4.**
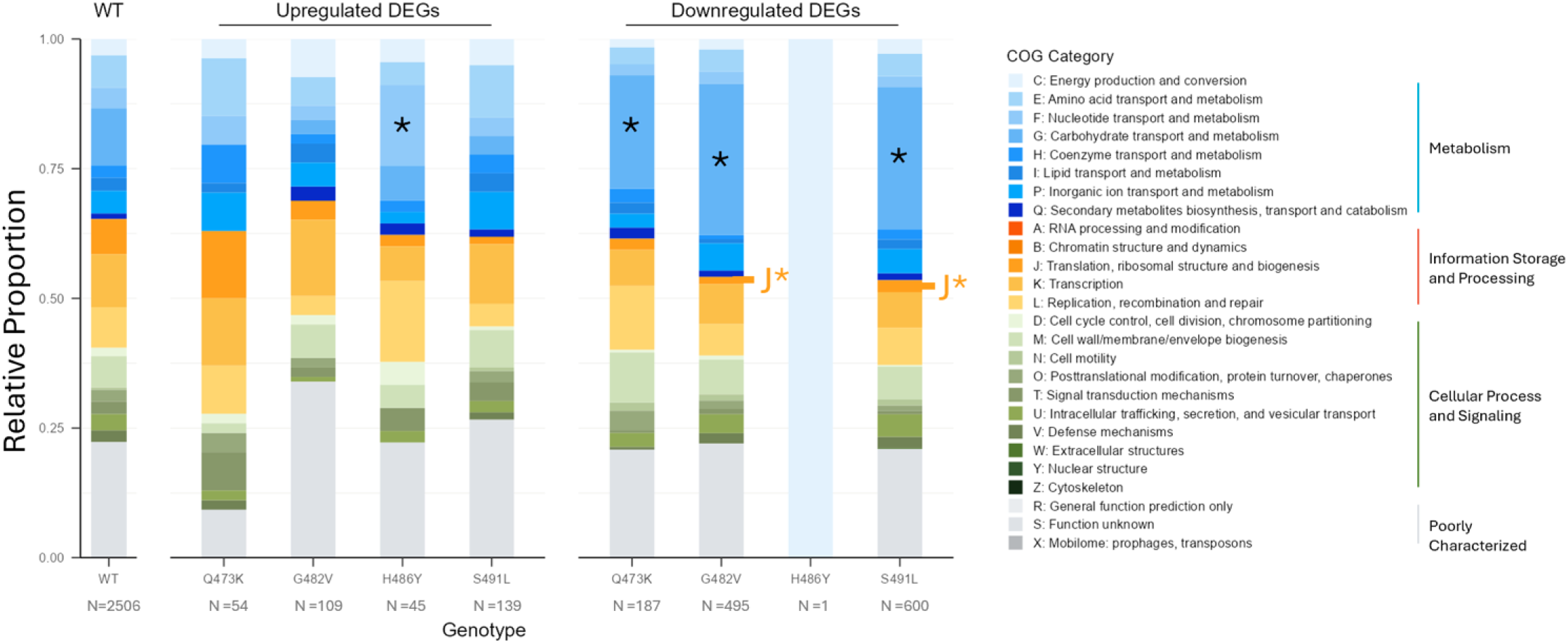
Distribution of COG categories in the wild-type (WT) strain genome and among differentially expressed genes in each RRDR mutant strain. COG categories were assigned using EggNOG-mapper and genes that did not have assigned COG categories were excluded. The distribution of differentially expressed genes in each RRDR mutant was compared with the distribution of all genes in the WT strain genome using Fisher’s exact test with Bonferroni correction. *p≤0.05.

### Allele-specific effects of RRDR mutations on *E. faecium* growth rate and antimicrobial susceptibility

Although RRDR mutations primarily decrease rifampin binding^18,28^, these mutations can also impact RNA synthesis and thus overall bacterial growth rates^20,29–32^. We assessed the impact of RRDR-mediated changes on *E. faecium* growth by comparing the lag phase duration and doubling time of each isogenic RRDR mutant strain with that of the WT strain. Among the four mutant strains, the H486Y mutant showed a clear growth advantage, as evidenced by a slightly shorter lag phase and faster doubling time (Figure 5A). The up-regulation of nucleotide transport and metabolism-associated genes in this mutant might contribute to this phenotype. In contrast, both the S491L and Q473K mutants showed significantly prolonged lag phases and doubling times, with the S491L mutant having the most severe growth defect of the four mutants tested. Interestingly, the G482V mutant displayed a mixed phenotype, with an extended lag phase but shorter doubling time relative to the WT strain (Figure 5A). These growth differences could be due to the significant down-regulation of carbohydrate transport and metabolism and translation-associated genes in these mutants (Figure 4; Supplemental File 1).

**Figure 5.**
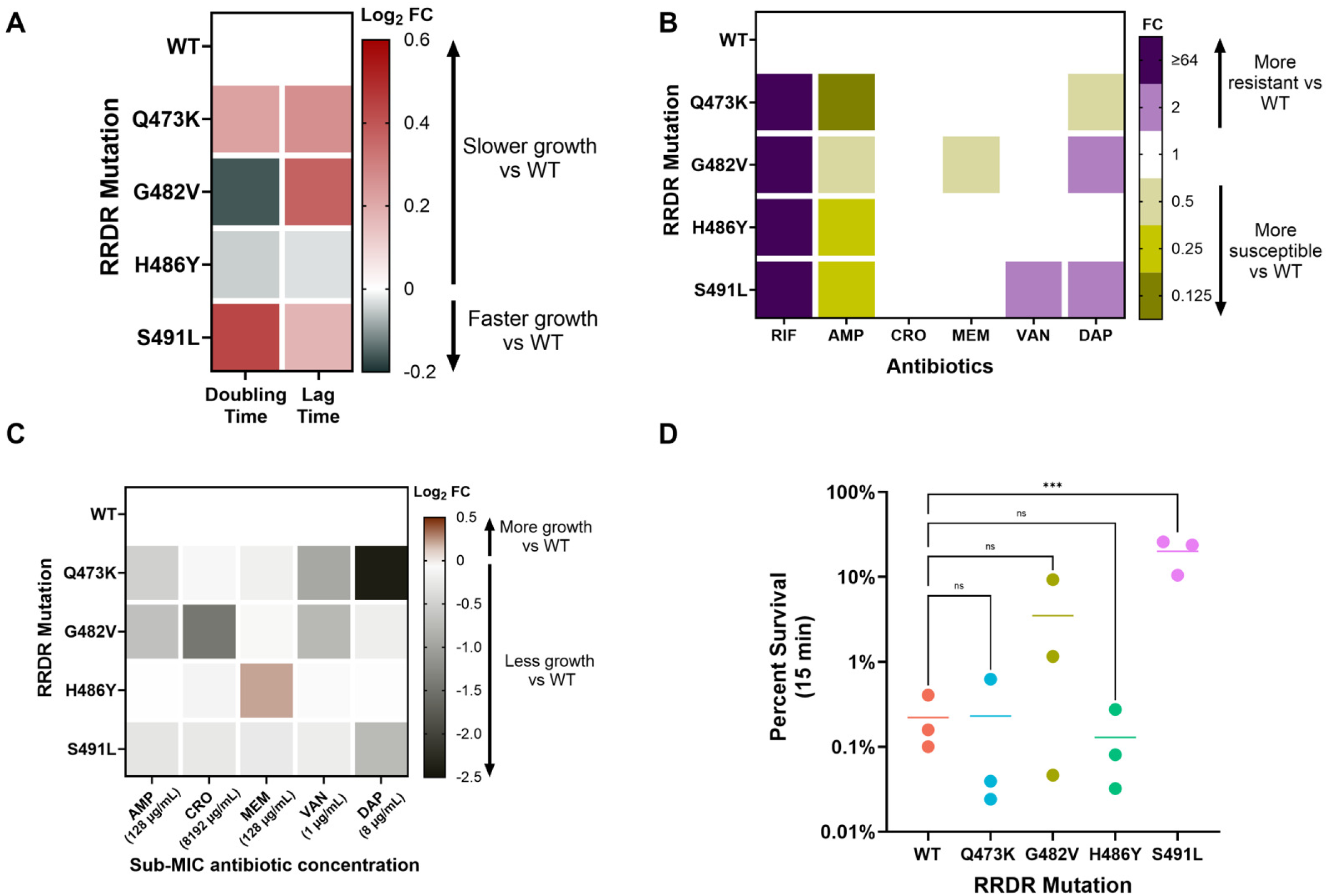
Allele-specific effects of RRDR mutations on *E. faecium* phenotypes. (A) Doubling and lag time of each strain were determined through growth assays in rich media. Results are shown as a heatmap of log_2_ fold change (FC) compared to the wild-type (WT) strain. (B) MICs as determined by broth microdilution. The MIC of each RRDR mutant is shown as FC compared to WT. (C) RRDR mutant growth under antibiotic stress, shown as the log_2_ FC of the total area under the growth curve (AUC) compared to WT. (D) Isopropanol tolerance displayed as percent survival after 15 min incubation with 20% isopropanol. All experiments were conducted in triplicate and mean values are shown. Significance was calculated with one-way ANOVA. *p≤0.05; **p≤0.01; ***p≤0.001; ****p≤0.0001; ns, not significant. RIF = Rifampin; AMP = Ampicillin; CRO = Ceftriaxone; MEM = Meropenem; VAN = Vancomycin; DAP = Daptomycin.

RRDR mutations have been known to influence susceptibility to antibiotics beyond those in the rifamycin class, in both *E. faecium* and other bacterial species^13,14,21,33,34^. To assess these effects, we measured the minimum inhibitory concentrations (MICs) of various antibiotics against the WT and RRDR mutant strains. As expected, all four mutants displayed increased rifampin resistance compared to the WT strain (Figure 5B). Additionally, all four mutants had decreased ampicillin MICs, ranging from 2-8-fold more susceptible. However, no differences in ceftriaxone susceptibility were noted, likely due to uniformly high MICs across all strains. Responses to meropenem and vancomycin were varied, with the G482V mutant displaying increased susceptibility to meropenem and the S491L mutant displaying increased resistance to vancomycin. Finally, the G482V and S491L mutant strains both had 2-fold higher MICs to daptomycin, while the Q473K mutant had a 2-fold lower daptomycin MIC compared to the WT strain. Together, these data suggest that RRDR mutations confer allele-specific effects on susceptibility to other antibiotics besides rifampin.

We also assessed differences in *E. faecium* fitness under antibiotic stress by determining the total area under the growth curve (AUC) in the presence of sub-MIC concentrations of the same antibiotics tested in Figure 5B. Consistent with the slower growth rates of the Q473K, G482V, and S491L strains, these mutants also displayed lower growth under sub-MIC antibiotic pressure (Figure 5C). The H486Y mutant, however, demonstrated better overall growth under meropenem stress, despite having the same MIC as the WT strain (Figure 5C). Moreover, the H486Y mutant showed no significant reduction in growth under any antibiotic stress compared to the WT strain (Figure 5C), perhaps due to the minimal transcriptional changes in this mutant (Figure 3C). Finally, we investigated whether RRDR mutations contribute to isopropanol tolerance^35^, and observed that the S491L mutant exhibited significantly higher tolerance to isopropanol compared to the WT strain (Figure 5D, Supplemental Figure 4C). Together these data suggest that the H486Y mutation and variation at residue 491 in the RRDR cause increased meropenem or isopropanol tolerance, either of which could facilitate the spread of *E. faecium* in clinical settings.

### Transcriptional changes in *E. faecium* carbohydrate metabolism genes are associated with differences in peptidoglycan abundance and muropeptide profiles

Given the changes we observed in mutant strain growth dynamics and expression of carbohydrate metabolism-associated genes, we wondered whether RRDR mutations also altered peptidoglycan abundance and composition. The H486Y mutant, which displayed minimal transcriptional changes and growth defects, produced similar amounts of peptidoglycan as the WT strain (Figure 6A). In contrast, the other three mutants, which all exhibited significant down-regulation of carbohydrate metabolism-associated genes, produced significantly less peptidoglycan compared to the WT strain (Figure 6A). The S491L mutant, which exhibited the most dramatic transcriptional changes and growth defects, produced the least peptidoglycan of all mutants (Figures 3D, 5A, and 6A). Together these data suggest that RRDR mutations alter *E. faecium* peptidoglycan abundance in an allele-specific manner.

**Figure 6.**
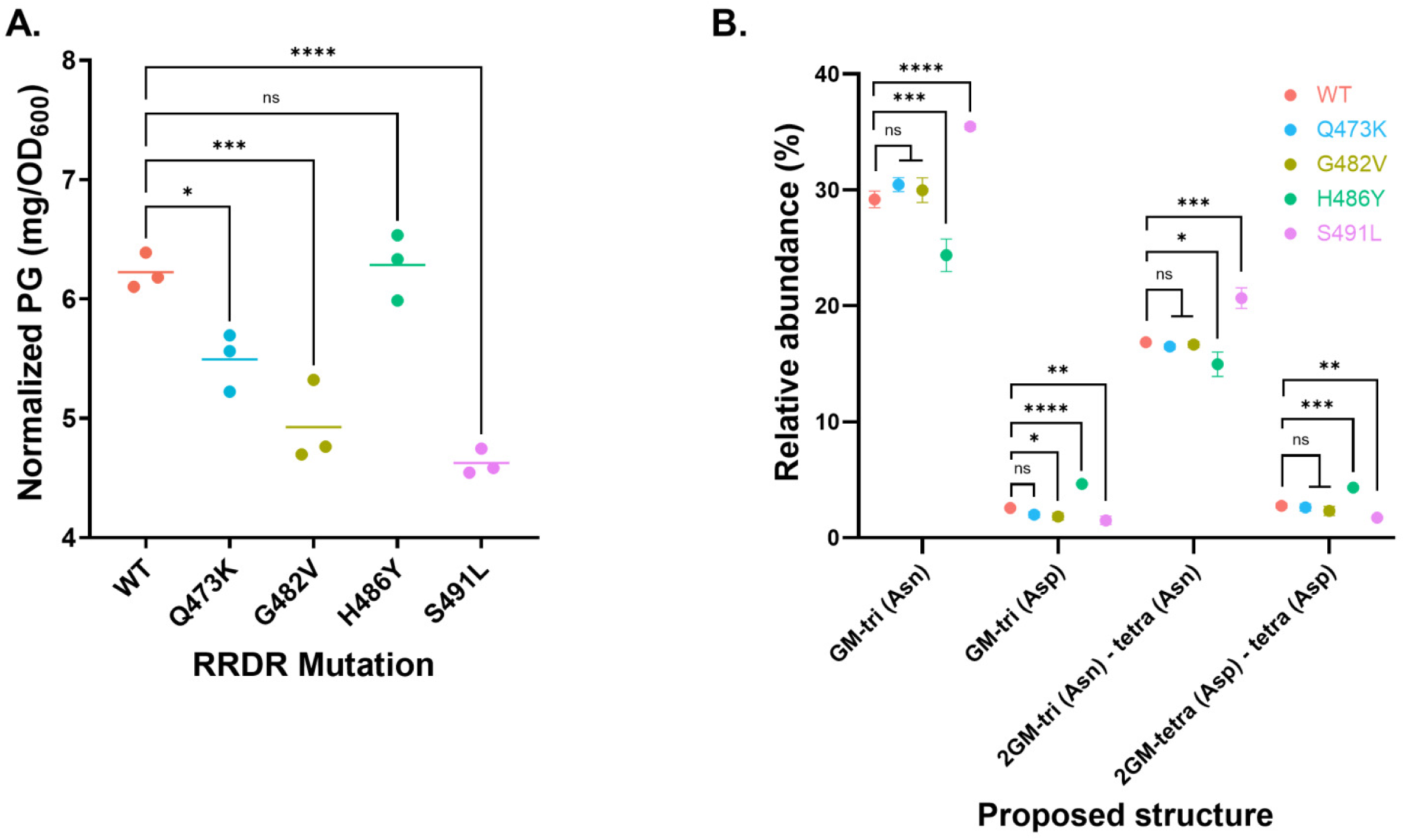
Changes in peptidoglycan abundance and composition due to RRDR mutation. (A) Normalized abundance of extracted peptidoglycan (PG) from wild type (WT) and mutant strains. (B) Relative abundance of muropeptides isolated from WT and mutant strains. GM, disaccharide (GlcNAc-MurNAc); 2GM, disaccharide-disaccharide (GlcNAc-MurNAc-GlcNAc-MurNAc); GM-Tri, disaccharide tripeptide (L-Ala-D-iGln-L-Lys); GM-Tetra, disaccharide tetrapeptide (L-Ala-D-iGln-L-Lys-D-Ala). Assignment of amide and hydroxyl functions to either peptide stem is arbitrary. Significance was calculated using one-way ANOVA with Tukey’s multiple comparison post-hoc test. *p≤0.05; **p≤0.01; ***p≤0.001; ****p≤0.0001; ns, not significant.

Further analysis of cell wall peptidoglycan fragments revealed significant differences in muropeptide composition between the WT and mutant *E. faecium* strains, particularly in amidation levels. The S491L mutant exhibited a significant increase in N-acetylglucosamine-muramyl tripeptide amidation compared to the WT strain (Figure 6B, Supplemental Figure 5A; increased GM-tri (Asn) and decreased GM-tri (Asp)). This heightened amidation could result from altered intracellular glutamine levels due to down-regulation of *nodM* (LKHMGDEB_00431) and up-regulation of *gadB* (LKHMGDEB_00941), which utilize L-glutamine and L-glutamate as substrates, respectively (Supplemental File 1)^36,37^. Increased amidation is often linked to greater peptidoglycan cross-linking, which can lead to higher metabolic cost, and thus slower rates of cell wall turnover and growth in the S491L mutant (Figure 5A and 6B; increased 2GM-tri (Asn) – tetra (Asn))^38^. Conversely, the muropeptide profile of the H486Y isolate showed significantly lower amidation levels. This may be associated with the up-regulation of *glmS* (LKHMGDEB_00994), which also consumes L-glutamine as a substrate, thereby decreasing glutamine availability in the cell and overall amidation levels and reducing the cost associated with cell wall synthesis^39^ (Figure 6B, Supplemental File 1). This reduced metabolic cost might contribute to the faster growth rate of the H486Y mutant compared to the WT strain (Figure 5A). While further studies are needed to fully understand the connection between RRDR mutations, transcriptional changes, alterations in cell wall composition, and changes in growth rate and antibiotic susceptibility, these data nonetheless highlight the variable effects of different RRDR mutations on *E. feacium* physiology.

## Discussion

To develop more effective anti-enterococcal therapeutics, a deeper understanding of how antimicrobial resistance mechanisms affect *E. faecium* physiology is critical. Here, we investigated the broader impacts of RRDR mutations beyond resistance to rifamycin antibiotics and their potential contribution to the global expansion of *E. faecium* as a hospital-adapted pathogen.

We found that *E. faecium* clinical isolates harboring RRDR mutations are prevalent and widely distributed both globally and locally. About 30% of global *E. faecium* genomes carried a mutation in the RRDR, with the most prevalent being S491F. The broad distribution of these mutations across various STs, countries, and genetic background as well as the diversity of variants found suggests that these mutations might confer significant advantages to the isolates harboring them. Further analyses of *E. faecium* clinical isolates from our local hospital showed that some RRDR mutations were associated with changes in susceptibility to daptomycin, a last-resort antibiotic used to treat vancomycin-resistant *E. faecium* infections. In contrast to a previous study^14^, we found that the S491F mutation was inversely associated with daptomycin resistance in our local dataset (Figure 2C). This discrepancy may be influenced by different genetic backgrounds, or by the fact that US isolates predominantly encode *vanA*-type vancomycin resistance while Australian isolates encode *vanB*^40^. Additionally, daptomycin MIC testing was performed on the Microscan platform, which might have overcalled resistance and could impact the associations we identified.

RRDR mutations have been previously shown to affect *E. faecium* bacterial transcription by altering nucleotide binding and/or protein stability^14^. In the isogenic mutants studied here, the G482V, H486Y, and S491L substitutions are predicted to introduce steric hindrance and disrupt hydrogen bonding in the RpoB active site, thereby affecting transcription efficiency. Meanwhile, the Q473K mutation replaces a neutral amino acid with a bulkier positively charged lysine, potentially causing electrostatic repulsion and altered RpoB activity. Transcriptomic and phenotypic analyses of these isogenic mutants revealed striking pleiotropic effects on *E. faecium* physiology. In general, RRDR mutations caused significant changes to *E. faecium* growth and gene expression profiles. Among the four mutants tested, the H486Y mutation stood out as the least disruptive, with a transcriptional profile most similar to WT. Moreover, this variant exhibited slight increases in growth rate both without and with meropenem pressure compared to the WT strain, perhaps due to increased expression of nucleotide transport and metabolism-associated genes as well as a lower metabolic cost due to decreased muropeptide cross-linking. The combination of minimal collateral costs and improved growth rates likely contributes to the prevalence of this variant in clinical settings, with over 5% of global *E. faecium* isolates encoding this mutation. The Q473K, G482V, and S491L mutant strains, on the other hand, all displayed significant transcriptional disruptions, with the S491L mutant having the largest number of differentially expressed genes. All three mutants showed significant down-regulation of genes involved in carbohydrate metabolism, which likely led to the attenuated growth rates we observed in these strains. Muropeptide profiling revealed a significant increase in amidation levels in the S491L mutant, suggesting a higher metabolic cost associated with cell wall remodeling that might further slow bacterial growth. The RNA-seq data pointed to increased intracellular glutamine levels as a possible driver of increased amidation. Notably, this mutant demonstrated increased tolerance to isopropanol, a selective advantage in the nosocomial settings where isopropanol-based disinfectants are widely used. Although S491L mutation is less prevalent than S491F, a prior study demonstrated that S491F produced phenotypic effects in *E. faecium* that are comparable to our findings with S491L, including a substantial fitness cost^14^. Given these parallels, it is plausible that S491F also confers increased isopropanol tolerance. Notably, the smaller number of genes with altered expression in the S491F mutant might explain the pervasiveness of this mutation rather than S491L in clinical settings as well as its enrichment in emerging *E. faecium* lineages ST80 and ST117^14,16^.

In addition to their effects on growth dynamics and transcription, we also explored whether RRDR mutations influenced other phenotypes important for *E. faecium* pathogenicity, such as antibiotic susceptibility. All four RRDR mutant strains were more resistant to rifampin, but they were also all more susceptible to ampicillin, although we did not find significant changes in the mutants’ penicillin-binding protein 5 expression level or sequence^41^. We were surprised to find that the S491L mutant had a higher vancomycin MIC compared with the WT strain. This mutant uniquely showed increased expression of a VanY-like D-alanyl-D-alanine carboxypeptidase, an enzyme involved in peptidoglycan remodeling and cell wall synthesis (Supplemental File 1)^42,43^. Although primarily associated with cell wall synthesis, the function of this gene suggests a mechanism similar to VanY, whereby removal of the terminal D-alanine from the peptidoglycan pentapeptide prevents vancomycin binding, potentially contributing a modest decrease in vancomycin susceptibility^43^.

We also determined the impact of RRDR mutations on susceptibility to daptomycin, a frequently prescribed antibiotic for vancomycin-resistant *E. faecium* infections. Daptomycin resistance can be achieved through various mechanisms, including modifications to cell membrane composition and/or cell surface charge^44–47^. Both the G482V and S491L mutants exhibited a 2-fold increase in daptomycin MIC; however, the mechanism by which this phenotype was likely achieved differed between the mutants. Although the G482V mutant exhibited upregulated expression of *dltAB,* which has been shown to increase positive cell surface charge and thus decrease daptomycin binding^48–50^, we did not observe increased positive charge in this mutant (Supplemental File 1, Supplemental Figure 4D). This discrepancy could be due to additional changes in carbohydrate metabolism-associated genes in this mutant (Figure 4). Regarding decreased daptomycin susceptibility of the S491L mutant, we suspect that increased muropeptide amidation might contribute to reduced daptomycin binding to the cell membrane^44,48,49^. While the H486Y mutant exhibited upregulation of genes within the daptomycin resistance-associated LiaFSR system, particularly *liaF, liaS, liaX,* and *liaY,* these changes were not sufficient to increase daptomycin resistance, an observation consistent with a prior study (Supplemental File 1)^46,48,51–53^. We also investigated the *prdRAB* operon, which was recently implicated in RpoB-mediated daptomycin resistance in *E. faecium*^14^. However, no significant changes in *prdRAB* transcription were detected among any of the mutant strains we studied (Supplementary File 1). This variation might reflect differences in genetic backgrounds or differences between the vancomycin-susceptible strains used here and vancomycin-resistant *E. faecium* strains tested previously.

This study had several limitations. First, we were unable to fully recapitulate the most prevalent mutants identified among global *E. faecium* isolates. Given the allele-specific changes we observed, it is unknown whether a different mutation at the same amino acid residue would cause similar effects on *E. faecium* physiology. Second, while each of the RRDR mutations we studied conferred distinct selective advantage *in vitro*, their role in *E. faecium* growth and survival during infection remain to be investigated. Finally, it is unclear how the presence of vancomycin resistance plasmids might influence the effects of RRDR mutations, especially since some of our findings were discrepant with those of a recent study^14^. Nonetheless, our findings are in agreement with prior reports showing that alterations in the RRDR lead to collateral impacts on *E. faecium* growth, gene expression, and antibiotic susceptibility^13,14,18,20,29,31,34,54^.

Overall, this study illustrates how different RRDR mutations conferring rifampin resistance have unique and allele-specific effects on *E. faecium* physiology, highlighting the complex interplay between RRDR mutations and transcriptional and phenotypic changes. Understanding how RRDR mutations emerge and spread can help inform novel targets for therapeutic strategies to mitigate *E. faecium* persistence and pathogenicity in hospital settings.

## Materials and Methods

### Global *E. faecium* isolate analysis

All *E. faecium* genomes deposited in NCBI between 2000 and 2023 were downloaded on March 26^th^, 2024. Genomes were filtered as described in Supplemental Figure 1. Briefly, only genomes with “geo_location”, “host”, and “isolation_source” entries were included. Samples were de-duplicated based on BioSample Accession, and only those for which “host” included *Homo sapiens, Homo sapiens sapiens,* or human were retained for further analysis. Genomes with a size of >2.6 Mbp and <3.3 Mbp (determined using QUAST v5.2.0^55^), and which were not isolated from the environment, were used for the final global dataset (total = 14,384 isolates). Multi-locus sequence types (STs) were identified using the PubMLST database^56^. RRDR variants were determined using Geneious v2024.0.3 to align the assemblies to a representative WT RRDR sequence (467-GSSQLSQFMDQTNPLGELTHKRRLSAL-493).

### Local *E. faecium* isolate analysis

A total of 710 vancomycin-resistant *E. faecium* isolates were collected between 2017 and 2022 as part of a retrospective observational study at the University of Pittsburgh Medical Center called the Enhanced Detection System for Healthcare-Associated Transmission (EDS-HAT)^17^. Prior antibiotic exposure data was assessed for the 90 days prior to isolate collection. Daptomycin susceptibilities were determined through automated testing on the MicroScan platform and were interpreted according to the 2025 Clinical and Laboratory Standards Institute M100 guidelines^57^. The study was approved by the Institutional Review Board at the University of Pittsburgh (STUDY21040126). Clinical isolate genomes were analyzed using the same pipeline as for global *E. faecium* isolates.

### Selection of RRDR mutant strains

For *in vitro* rifampicin resistance selection, a previously described ST203 *E. faecium* strain called DVT705 was used as the parent wild-type (WT) strain, with antibiotic susceptibilities summarized in Supplementary File 1^16^. To select isogenic strains with RRDR mutations, an overnight culture of DVT705 was plated onto several brain heart infusion (BHI) agar plates containing 50 µg/mL rifampin. At least three different colonies were picked from each plate and the RRDR region was Sanger-sequenced. Four isolates with unique mutations were identified and further characterized. All *E. faecium* strains were grown in BHI broth at 37°C with agitation at 225 rpm. To confirm the isogenic mutant strains with whole genome sequencing, genomic DNA was extracted using a DNeasy Blood and Tissue Kit (Qiagen, Germantown, MD) according to the manufacturer’s protocol with additional incubation with 10 µL of 2500 U/mL mutanolysin and 50 µL 50 mg/mL lysozyme at 37°C for 1 hr. Next-generation sequencing libraries were prepared (2x150 bp, paired-end reads) and sequenced on the Illumina platform at SeqCenter (Pittsburgh, PA). The resulting reads were assembled using SPAdes v3.15.5 and annotated with Prokka v1.14.5^58,59^. Variants in each isogenic mutant were confirmed using breseq v0.37.1^60^.

### *E. faecium* growth dynamics and antimicrobial susceptibility testing

Changes in growth rates and antibiotic minimum inhibitory concentrations (MICs) of WT and isogenic mutant strains were determined as follows. Briefly, overnight cultures were normalized to OD_600_ of 0.05, and 5 µL of the diluted culture was used to inoculate 200 uL fresh BHI media in 96-well microplates (1:40 dilution). OD_600_ readings were taken every 20 minutes at 37°C for 24 hours using a Synergy H1 microplate reader (Biotek, Winooski, VT). For lag time calculations, the start of log-phase growth was set to a threshold of OD_600_ = 0.1. Antibiotic MICs were determined using the broth microdilution method^57^. Daptomycin MICs were determined with the addition of 50 µg/mL CaCl_2_ to the media. To assess growth in the presence of sub-MIC antibiotic concentrations, the following were tested: ampicillin at 128 µg/mL, ceftriaxone at 8192 µg/mL, meropenem at 128 µg/mL, vancomycin at 1 µg/mL, and daptomycin at 8 µg/mL. Results are presented as heatmaps showing log_2_ fold changes relative to the WT strain.

Isopropanol tolerance was determined as previously described^35^. Briefly, overnight cultures were diluted to OD_600_ of 1.66 in fresh BHI. Either 1 mL of phosphate-buffered saline (PBS; positive control) or isopropanol at 20% final concentration was added to 300 µL of the diluted cultures. Samples were vortexed for 5 seconds and incubated at room temperature for 5 or 15 minutes without shaking. Just prior to dilution, samples were vortexed again for 5 seconds. Serial dilutions (10^-1^ to 10^-7^) were prepared in PBS containing 7.5% Tween 80 and were spotted onto BHI plates to enumerate the colony forming units per mL (CFU/mL). Plates were incubated overnight at 37°C and CFU/mL were compared between PBS-treated and isopropanol-treated samples. Percent survival was calculated as the ratio of CFU/mL of isopropanol-treated samples to CFU/mL of PBS-treated samples. All experiments were conducted in biological triplicate.

### RNA sequencing and analysis

RNA was extracted from WT and isogenic mutant strains using a RNeasy Mini Kit (Qiagen) according to the manufacturer’s protocol with minor modifications. Briefly, 2x v/v of RNAprotect (Qiagen) was added to 5 mL mid-log phase cultures and samples were pelleted. The pellet was incubated with 10 µL of 2500 U/mL mutanolysin and 50 µL 50 mg/mL lysozyme at 37°C for 30 minutes prior to RNA extraction. Total RNA was treated with DNase and 2x150 bp paired-end libraries were generated and sequenced on an Illumina NovaSeq X Plus (SeqCenter, Pittsburgh, PA). Four replicate samples were collected for each strain. Sequencing reads were processed using Trim galore v0.6.10^61^. Reads passed quality control if the Phred score was >33 and the sequence length was >142 bp. Trimmed reads were then mapped to the annotated WT genome using STAR v2.7.11b^62^. Reads mapping to coding sequences were counted using featureCounts v2.0.6 against the annotated WT sequence^63^. Differential gene expression analysis was conducted using the DESeq2 package in RStudio^64^. For reliability, only genes with read counts >10 were included in the analysis. Genes were considered significantly differentially expressed if they had an adjusted *P*-value <0.05 (Benjamini-Hochberg for false discovery rate) and log_2_ fold change values <-1 or >1. Data were normalized with the varianceStabilizingTransformation() function for Principal Component Analysis (PCA). Volcano plots and Venn diagram were constructed using the EnhancedVolcano and VennDiagram packages in RStudio, respectively.

The Clusters of Orthologous Groups (COG) database was used to assign functions to differentially expressed genes using EggNOG-mapper^26,27,65^. Approximately 84% (2388/2854) of annotated genes in the WT strain were assigned to a COG category. For distribution analysis, genes assigned to multiple COG categories were counted once in each category to yield a total of 2,506 categorized genes. DEGs without a specified COG category were excluded from analysis.

### Peptidoglycan LC-MS analysis

Peptidoglycan was extracted from the sacculi of WT and isogenic mutant strains and digested with 10 KU/mL mutanolysin as previously described^66^. For LC-MS analysis, the peptidoglycan of each strain was further analyzed as previously described^67^. Briefly, peptidoglycan was digested with mutanolysin and then treated with sodium borohydride in 0.25 M boric acid at pH 9 for 1 hour at room temperature. Orthophosphoric acid was then used to quench the reaction and the pH was adjusted to 2 – 3. After centrifugation at 20,000xg for 10 minutes, the peptidoglycan was then analyzed on a 1290 Infinity II LC/MSD system (Agilent technologies) using a Poroshell 120 EC-C18 column. The mobile phase consisted of 0.1% formic acid in water and the eluent was 0.1% formic acid in acetonitrile. Samples were first run against the mobile phase at a flow rate of 0.5 mL/ min, then against 2% (0 – 5 min) and 2 – 10% (5 – 65 min) eluent. Absorbance of the eluting peaks was measured at 205 nm and detected with MSD API-ES Scan mode (m/z = 200 – 2500). Area under the curve of each individual peak from the chromatogram was calculated, and the relative abundance of each muropeptide was calculated as the percentage of each individual peak relative to all assigned peaks. The experiment was performed on three biological replicates.

### Cytochrome C binding assay

Cytochrome C binding was performed on WT and isogenic mutants as previously described with minor modifications^68^. Briefly, overnight cultures were washed three times with 1 mL of 20 mM 3-(N-Morpholino) propanesulfonic acid sodium salt buffer (MOPS, pH = 7.0) and normalized to OD_600_ = 0.2. The normalized suspension was then incubated with an equal volume of 1 mg/mL cytochrome C (Sigma Aldrich). After 30 minutes incubation at room temperature, samples were centrifuged for 5 minutes at 13,000 rpm. The absorbance of the supernatant was measured at 410 nm in a Synergy H1 microplate reader (Biotek, Winooski, VT) and results were averaged across technical triplicates. This experiment was conducted in biological triplicate.

### Statistical Analysis

Significance for COG category enrichment and associations between RRDR mutations with prior rifamycin class exposure and daptomycin susceptibility was assessed using Fisher’s exact test, with Bonferroni-correction when appropriate. Significance for growth rate, isopropanol tolerance, peptidoglycan analysis, and cytochrome c binding assays were assessed using one-way ANOVA, with Tukey’s multiple comparison post-hoc test when appropriate. All calculations were conducted in GraphPad Prism (v.10.0).

## Data availability

Genomes from the global collection are listed in Supplemental File 1. Genomes from the local collection are deposited in NCBI under BioProject PRJNA475751. RNA-sequencing data is deposited in the Gene Expression Omnibus (GEO) at NCBI under accession GSE302807.

## Acknowledgments

This work was supported by National Institute of Allergy and Diseases (R01 AI165519 to DVT) and by the Department of Medicine at the University of Pittsburgh School of Medicine. We gratefully acknowledge Alexander Sundermann, Lee Harrison, and Lora Pless for providing access to isolate genomes collected through the EDS-HAT program (R01 AI127472), as well as Lloyd Clarke for assistance collecting antibiotic exposure data. The funders had no role in study design, data collection and analysis, decision to publish, or preparation of the manuscript.

## References

1. Naghavi, M., Vollset, S.E., Ikuta, K.S., Swetschinski, L.R., Gray, A.P., Wool, E.E., Aguilar, G.R., Mestrovic, T., Smith, G., Han, C., et al. (2024). Global burden of bacterial antimicrobial resistance 1990–2021: a systematic analysis with forecasts to 2050. The Lancet 404, 1199–1226. 10.1016/S0140-6736(24)01867-1.

2. O’Neill, J. (2016). Tackling drug-resistant infections globally: final report and recommendations (Government of the United Kingdom).

3. Ubeda, C., Taur, Y., Jenq, R.R., Equinda, M.J., Son, T., Samstein, M., Viale, A., Socci, N.D., van den Brink, M.R.M., Kamboj, M., et al. (2010). Vancomycin-resistant Enterococcus domination of intestinal microbiota is enabled by antibiotic treatment in mice and precedes bloodstream invasion in humans. J. Clin. Invest. 120, 4332–4341. 10.1172/JCI43918.

4. Rice, L.B. (2008). Federal Funding for the Study of Antimicrobial Resistance in Nosocomial Pathogens: No ESKAPE. J. Infect. Dis. 197, 1079–1081. 10.1086/533452.

5. Arias, C.A., and Miller, W.R. (2023). Mechanisms of antibiotic resistance in enterococci. https://www.uptodate.com/contents/mechanisms-of-antibiotic-resistance-in-enterococci.

6. 6. Van Tyne, D., and Gilmore, M.S. (2014). Friend Turned Foe: Evolution of Enterococcal Virulence and Antibiotic Resistance. Annu. Rev. Microbiol. 68, 337–356. 10.1146/annurev-micro-091213-113003.

7. Arias, C.A., and Murray, B.E. (2012). The rise of the Enterococcus: beyond vancomycin resistance. Nat. Rev. Microbiol. 10, 266–278. 10.1038/nrmicro2761.

8. Chen, Q., Yin, D., Li, P., Guo, Y., Ming, D., Lin, Y., Yan, X., Zhang, Z., and Hu, F. (2020). First Report Cfr and OptrA Co-harboring Linezolid-Resistant Enterococcus faecalis in China. Infect. Drug Resist. 13, 3919–3922. 10.2147/IDR.S270701.

9. Turnidge, J., Kahlmeter, G., Cantón, R., MacGowan, A., and Giske, C.G. (2020). Daptomycin in the treatment of enterococcal bloodstream infections and endocarditis: a EUCAST position paper. Clin. Microbiol. Infect. 26, 1039–1043. 10.1016/j.cmi.2020.04.027.

10. Dadashi, M., Sharifian, P., Bostanshirin, N., Hajikhani, B., Bostanghadiri, N., Khosravi-Dehaghi, N., van Belkum, A., and Darban-Sarokhalil, D. (2021). The Global Prevalence of Daptomycin, Tigecycline, and Linezolid-Resistant Enterococcus faecalis and Enterococcus faecium Strains From Human Clinical Samples: A Systematic Review and Meta-Analysis. Front. Med. 8, 720647. 10.3389/fmed.2021.720647.

11. Gilmore, M.S., Lebreton, F., and van Schaik, W. (2013). Genomic Transition of Enterococci from Gut Commensals to Leading Causes of Multidrug-resistant Hospital Infection in the Antibiotic Era. Curr. Opin. Microbiol. 16, 10–16. 10.1016/j.mib.2013.01.006.

12. Murray, C.J., Ikuta, K.S., Sharara, F., Swetschinski, L., Aguilar, G.R., Gray, A., Han, C., Bisignano, C., Rao, P., Wool, E., et al. (2022). Global burden of bacterial antimicrobial resistance in 2019: a systematic analysis. The Lancet 399, 629–655. 10.1016/S0140-6736(21)02724-0.

13. Kristich, C.J., and Little, J.L. (2012). Mutations in the β Subunit of RNA Polymerase Alter Intrinsic Cephalosporin Resistance in Enterococci. Antimicrob. Agents Chemother. 56, 2022–2027. 10.1128/AAC.06077-11.

14. Turner, A.M., Li, L., Monk, I.R., Lee, J.Y.H., Ingle, D.J., Portelli, S., Sherry, N.L., Isles, N., Seemann, T., Sharkey, L.K., et al. (2024). Rifaximin prophylaxis causes resistance to the last-resort antibiotic daptomycin. Nature 635, 969–977. 10.1038/s41586-024-08095-4.

15. Lane, W.J., and Darst, S.A. (2010). Molecular Evolution of Multisubunit RNA Polymerases: Sequence Analysis. J. Mol. Biol. 395, 671–685. 10.1016/j.jmb.2009.10.062.

16. Mills, E.G., Hewlett, K., Smith, A.B., Griffith, M.P., Pless, L., Sundermann, A.J., Harrison, L.H., Zackular, J.P., and Van Tyne, D. (2025). Bacteriocin production facilitates nosocomial emergence of vancomycin-resistant Enterococcus faecium. Nat. Microbiol., 1–11. 10.1038/s41564-025-01958-0.

17. Sundermann, A.J., Chen, J., Kumar, P., Ayres, A.M., Cho, S.T., Ezeonwuka, C., Griffith, M.P., Miller, J.K., Mustapha, M.M., Pasculle, A.W., et al. (2022). Whole-Genome Sequencing Surveillance and Machine Learning of the Electronic Health Record for Enhanced Healthcare Outbreak Detection. Clin. Infect. Dis. 75, 476–482. 10.1093/cid/ciab946.

18. Molodtsov, V., Scharf, N.T., Stefan, M.A., Garcia, G.A., and Murakami, K.S. (2017). Structural basis for rifamycin resistance of bacterial RNA polymerase by the three most clinically important RpoB mutations found in Mycobacterium tuberculosis. Mol. Microbiol. 103, 1034–1045. 10.1111/mmi.13606.

19. Goldstein, B.P. (2014). Resistance to rifampicin: a review. J. Antibiot. (Tokyo) 67, 625–630. 10.1038/ja.2014.107.

20. Cutugno, L., Mc Cafferty, J., Pané-Farré, J., O’Byrne, C., and Boyd, A. (2020). rpoB mutations conferring rifampicin-resistance affect growth, stress response and motility in Vibrio vulnificus. Microbiology 166, 1160–1170. 10.1099/mic.0.000991.

21. Palace, S.G., Wang, Y., Rubin, D.H., Welsh, M.A., Mortimer, T.D., Cole, K., Eyre, D.W., Walker, S., and Grad, Y.H. (2020). RNA polymerase mutations cause cephalosporin resistance in clinical Neisseria gonorrhoeae isolates. eLife 9, e51407. 10.7554/eLife.51407.

22. Matsuo, M., Hishinuma, T., Katayama, Y., Cui, L., Kapi, M., and Hiramatsu, K. (2011). Mutation of RNA Polymerase β Subunit (rpoB) Promotes hVISA-to-VISA Phenotypic Conversion of Strain Mu3▿. Antimicrob. Agents Chemother. 55, 4188–4195. 10.1128/AAC.00398-11.

23. Perkins, A.E., and Nicholson, W.L. (2008). Uncovering New Metabolic Capabilities of Bacillus subtilis Using Phenotype Profiling of Rifampin-Resistant rpoB Mutants. J. Bacteriol. 190, 807–814. 10.1128/jb.00901-07.

24. Hall, A.R. (2013). Genotype-by-environment interactions due to antibiotic resistance and adaptation in Escherichia coli. J. Evol. Biol. 26, 1655–1664. 10.1111/jeb.12172.

25. Chilambi, G.S., Nordstrom, H.R., Evans, D.R., Ferrolino, J.A., Hayden, R.T., Marón, G.M., Vo, A.N., Gilmore, M.S., Wolf, J., Rosch, J.W., et al. (2020). Evolution of vancomycin-resistant Enterococcus faecium during colonization and infection in immunocompromised pediatric patients. Proc. Natl. Acad. Sci. 117, 11703–11714. 10.1073/pnas.1917130117.

26. Cantalapiedra, C.P., Hernández-Plaza, A., Letunic, I., Bork, P., and Huerta-Cepas, J. (2021). eggNOG-mapper v2: Functional Annotation, Orthology Assignments, and Domain Prediction at the Metagenomic Scale. Mol. Biol. Evol. 38, 5825–5829. 10.1093/molbev/msab293.

27. Galperin, M.Y., Wolf, Y.I., Makarova, K.S., Vera Alvarez, R., Landsman, D., and Koonin, E.V. (2021). COG database update: focus on microbial diversity, model organisms, and widespread pathogens. Nucleic Acids Res. 49, D274–D281. 10.1093/nar/gkaa1018.

28. Gill, S.K., and Garcia, G.A. (2011). Rifamycin inhibition of WT and Rif-resistant Mycobacterium tuberculosis and Escherichia coli RNA polymerases in vitro. Tuberculosis 91, 361–369. 10.1016/j.tube.2011.05.002.

29. Enne, V.I., Delsol, A.A., Roe, J.M., and Bennett, P.M. (2004). Rifampicin resistance and its fitness cost in Enterococcus faecium. J. Antimicrob. Chemother. 53, 203–207. 10.1093/jac/dkh044.

30. Reynolds, M.G. (2000). Compensatory Evolution in Rifampin-Resistant Escherichia coli. Genetics 156, 1471–1481. 10.1093/genetics/156.4.1471.

31. Wichelhaus, T.A., Böddinghaus, B., Besier, S., Schäfer, V., Brade, V., and Ludwig, A. (2002). Biological Cost of Rifampin Resistance from the Perspective of Staphylococcus aureus. Antimicrob. Agents Chemother. 46, 3381–3385. 10.1128/AAC.46.11.3381-3385.2002.

32. Qi, Q., Preston, G.M., and MacLean, R.C. (2014). Linking System-Wide Impacts of RNA Polymerase Mutations to the Fitness Cost of Rifampin Resistance in Pseudomonas aeruginosa. mBio 5, 10.1128/mbio.01562-14. 10.1128/mbio.01562-14.

33. Friedman, L., Alder, J.D., and Silverman, J.A. (2006). Genetic Changes That Correlate with Reduced Susceptibility to Daptomycin in Staphylococcus aureus. Antimicrob. Agents Chemother. 50, 2137–2145. 10.1128/aac.00039-06.

34. Cui, L., Isii, T., Fukuda, M., Ochiai, T., Neoh, H., Camargo, I.L.B. da C., Watanabe, Y., Shoji, M., Hishinuma, T., and Hiramatsu, K. (2010). An RpoB Mutation Confers Dual Heteroresistance to Daptomycin and Vancomycin in Staphylococcus aureus. Antimicrob. Agents Chemother. 54, 5222–5233. 10.1128/AAC.00437-10.

35. Pidot, S.J., Gao, W., Buultjens, A.H., Monk, I.R., Guerillot, R., Carter, G.P., Lee, J.Y.H., Lam, M.M.C., Grayson, M.L., Ballard, S.A., et al. (2018). Increasing tolerance of hospital Enterococcus faecium to handwash alcohols. Sci. Transl. Med. 10, eaar6115. 10.1126/scitranslmed.aar6115.

36. Ghosh, S., Blumenthal, H.J., Davidson, E., and Roseman, S. (1960). Glucosamine Metabolism: V. ENZYMATIC SYNTHESIS OF GLUCOSAMINE 6-PHOSPHATE. J. Biol. Chem. 235, 1265–1273. 10.1016/S0021-9258(18)69397-4.

37. De Biase, D., Tramonti, A., Bossa, F., and Visca, P. (1999). The response to stationary-phase stress conditions in Escherichia coli: role and regulation of the glutamic acid decarboxylase system. Mol. Microbiol. 32, 1198–1211. 10.1046/j.1365-2958.1999.01430.x.

38. Apostolos, A.J., and Pires, M.M. (2022). Chapter Eleven - Impact of crossbridge structure on peptidoglycan crosslinking: A synthetic stem peptide approach. In Methods in Enzymology Chemical Microbiology Part B., E. E. Carlson, ed. (Academic Press), pp. 259–279. 10.1016/bs.mie.2021.11.019.

39. Cui, L., Murakami, H., Kuwahara-Arai, K., Hanaki, H., and Hiramatsu, K. (2000). Contribution of a Thickened Cell Wall and Its Glutamine Nonamidated Component to the Vancomycin Resistance Expressed by Staphylococcus aureus Mu50. Antimicrob. Agents Chemother. 44, 2276–2285. 10.1128/aac.44.9.2276-2285.2000.

40. Hughes, A., Ballard, S., Sullivan, S., and Marshall, C. (2019). An outbreak of vanA vancomycin-resistant Enterococcus faecium in a hospital with endemic vanB VRE. Infect. Dis. Health 24, 82–91. 10.1016/j.idh.2018.12.002.

41. Montealegre, M.C., Roh, J.H., Rae, M., Davlieva, M.G., Singh, K.V., Shamoo, Y., and Murray, B.E. (2016). Differential Penicillin-Binding Protein 5 (PBP5) Levels in the Enterococcus faecium Clades with Different Levels of Ampicillin Resistance. Antimicrob. Agents Chemother. 61, e02034–16. 10.1128/AAC.02034-16.

42. Ghosh, A.S., Chowdhury, C., and Nelson, D.E. (2008). Physiological functions of D-alanine carboxypeptidases in Escherichia coli. Trends Microbiol. 16, 309–317. 10.1016/j.tim.2008.04.006.

43. Arthur, M., Depardieu, F., Snaith, H.A., Reynolds, P.E., and Courvalin, P. (1994). Contribution of VanY D,D-carboxypeptidase to glycopeptide resistance in Enterococcus faecalis by hydrolysis of peptidoglycan precursors. Antimicrob. Agents Chemother. 38, 1899–1903. 10.1128/aac.38.9.1899.

44. Supandy, A., Mehta, H.H., Tran, T.T., Miller, W.R., Zhang, R., Xu, L., Arias, C.A., and Shamoo, Y. (2022). Evolution of Enterococcus faecium in Response to a Combination of Daptomycin and Fosfomycin Reveals Distinct and Diverse Adaptive Strategies. Antimicrob. Agents Chemother. 66, e02333–21. 10.1128/aac.02333-21.

45. Diaz, L., Tran, T.T., Munita, J.M., Miller, W.R., Rincon, S., Carvajal, L.P., Wollam, A., Reyes, J., Panesso, D., Rojas, N.L., et al. (2014). Whole-Genome Analyses of Enterococcus faecium Isolates with Diverse Daptomycin MICs. Antimicrob. Agents Chemother. 58, 4527–4534. 10.1128/AAC.02686-14.

46. Prater, A.G., Mehta, H.H., Kosgei, A.J., Miller, W.R., Tran, T.T., Arias, C.A., and Shamoo, Y. (2019). Environment Shapes the Accessible Daptomycin Resistance Mechanisms in Enterococcus faecium. Antimicrob. Agents Chemother., AAC.00790–19. 10.1128/AAC.00790-19.

47. Tran, T.T., Munita, J.M., and Arias, C.A. (2015). Mechanisms of Drug Resistance: Daptomycin Resistance. Ann. N. Y. Acad. Sci. 1354, 32–53. 10.1111/nyas.12948.

48. Prater, A.G., Mehta, H.H., Beabout, K., Supandy, A., Miller, W.R., Tran, T.T., Arias, C.A., and Shamoo, Y. (2021). Daptomycin resistance in Enterococcus faecium can be delayed by disruption of the LiaFSR stress response pathway. Antimicrob. Agents Chemother. 10.1128/AAC.01317-20.

49. Grein, F., Müller, A., Scherer, K.M., Liu, X., Ludwig, K.C., Klöckner, A., Strach, M., Sahl, H.-G., Kubitscheck, U., and Schneider, T. (2020). Ca 2+ -Daptomycin targets cell wall biosynthesis by forming a tripartite complex with undecaprenyl-coupled intermediates and membrane lipids. Nat. Commun. 11, 1–11. 10.1038/s41467-020-15257-1.

50. Yang, S.-J., Kreiswirth, B.N., Sakoulas, G., Yeaman, M.R., Xiong, Y.Q., Sawa, A., and Bayer, A.S. (2009). Enhanced expression of dltABCD is associated with development of daptomycin nonsusceptibility in a clinical endocarditis isolate of Staphylococcus aureus. J. Infect. Dis. 200, 1916–1920. 10.1086/648473.

51. Panesso, D., Reyes, J., Gaston, E.P., Deal, M., Londoño, A., Nigo, M., Munita, J.M., Miller, W.R., Shamoo, Y., Tran, T.T., et al. (2015). Deletion of liaR Reverses Daptomycin Resistance in Enterococcus faecium Independent of the Genetic Background. Antimicrob. Agents Chemother. 59, 7327–7334. 10.1128/AAC.01073-15.

52. Nguyen, A.H., Tran, T.T., Panesso, D., Hood, K.S., Polamraju, V., Zhang, R., Khan, A., Miller, W.R., Mileykovskaya, E., Shamoo, Y., et al. (2024). Molecular basis of cell membrane adaptation in daptomycin-resistant Enterococcus faecalis. JCI Insight 9, e173836. 10.1172/jci.insight.173836.

53. Miller, C., Kong, J., Tran, T.T., Arias, C.A., Saxer, G., and Shamoo, Y. (2013). Adaptation of Enterococcus faecalis to Daptomycin Reveals an Ordered Progression to Resistance. Antimicrob. Agents Chemother. 57, 5373–5383. 10.1128/AAC.01473-13.

54. Patel, Y., Soni, V., Rhee, K.Y., and Helmann, J.D. (2023). Mutations in rpoB That Confer Rifampicin Resistance Can Alter Levels of Peptidoglycan Precursors and Affect β-Lactam Susceptibility. mBio 14, e03168–22. 10.1128/mbio.03168-22.

55. Gurevich, A., Saveliev, V., Vyahhi, N., and Tesler, G. (2013). QUAST: quality assessment tool for genome assemblies. Bioinformatics 29, 1072–1075. 10.1093/bioinformatics/btt086.

56. Jolley, K.A., Bray, J.E., and Maiden, M.C.J. (2018). Open-access bacterial population genomics: BIGSdb software, the PubMLST.org website and their applications. Wellcome Open Res. 3, 124. 10.12688/wellcomeopenres.14826.1.

57. CLSI (2025). Performance Standards for Antimicrobial Susceptibility Testing, 35th ed. (Clinical and Laboratory Standards Institute).

58. Bankevich, A., Nurk, S., Antipov, D., Gurevich, A.A., Dvorkin, M., Kulikov, A.S., Lesin, V.M., Nikolenko, S.I., Pham, S., Prjibelski, A.D., et al. (2012). SPAdes: A New Genome Assembly Algorithm and Its Applications to Single-Cell Sequencing. J. Comput. Biol. 19, 455–477. 10.1089/cmb.2012.0021.

59. Seemann, T. (2014). Prokka: rapid prokaryotic genome annotation. Bioinformatics 30, 2068–2069. 10.1093/bioinformatics/btu153.

60. Deatherage, D.E., and Barrick, J.E. (2014). Identification of mutations in laboratory evolved microbes from next-generation sequencing data using breseq. Methods Mol. Biol. Clifton NJ 1151, 165–188. 10.1007/978-1-4939-0554-6_12.

61. Krueger, F. (2021). TrimGalore. https://github.com/FelixKrueger/TrimGalore

62. Dobin, A., Davis, C.A., Schlesinger, F., Drenkow, J., Zaleski, C., Jha, S., Batut, P., Chaisson, M., and Gingeras, T.R. (2013). STAR: ultrafast universal RNA-seq aligner. Bioinformatics 29, 15–21. 10.1093/bioinformatics/bts635.

63. Liao, Y., Smyth, G.K., and Shi, W. (2014). featureCounts: an efficient general purpose program for assigning sequence reads to genomic features. Bioinformatics 30, 923–930. 10.1093/bioinformatics/btt656.

64. Love, M.I., Huber, W., and Anders, S. (2014). Moderated estimation of fold change and dispersion for RNA-seq data with DESeq2. Genome Biol. 15, 550. 10.1186/s13059-014-0550-8.

65. Tatusov, R.L., Galperin, M.Y., Natale, D.A., and Koonin, E.V. (2000). The COG database: a tool for genome-scale analysis of protein functions and evolution. Nucleic Acids Res. 28, 33–36.

66. Kim, B., Wang, Y.-C., Hespen, C.W., Espinosa, J., Salje, J., Rangan, K.J., Oren, D.A., Kang, J.Y., Pedicord, V.A., and Hang, H.C. (2019). Enterococcus faecium secreted antigen A generates muropeptides to enhance host immunity and limit bacterial pathogenesis. eLife 8, e45343. 10.7554/eLife.45343.

67. Klupt, S., Fam, K.T., Zhang, X., Chodisetti, P.K., Mehmood, A., Boyd, T., Grotjahn, D., Park, D., and Hang, H.C. (2024). Secreted antigen A peptidoglycan hydrolase is essential for Enterococcus faecium cell separation and priming of immune checkpoint inhibitor therapy. eLife 13, RP95297. 10.7554/eLife.95297.

68. Miller, W.R., Nguyen, A., Singh, K.V., Rizvi, S., Khan, A., Erickson, S.G., Egge, S.L., Cruz, M., Dinh, A.Q., Diaz, L., et al. (2024). Membrane Lipids Augment Cell Envelope Stress Signaling and Resistance to Antibiotics and Antimicrobial Peptides in Enterococcus faecalis. J. Infect. Dis. 231, 307–317. 10.1093/infdis/jiae173.

